# Improved reference genome of the arboviral vector *Aedes albopictus*

**DOI:** 10.1101/2020.02.28.969527

**Authors:** U. Palatini, R.A. Masri, L.V. Cosme, S. Koren, F. Thibaud-Nissen, J.K. Biedler, F. Krsticevic, J.S. Johnston, R. Halbach, J.E. Crawford, I. Antoshechkin, A. Failloux, E. Pischedda, M. Marconcini, J. Ghurye, A. Rhie, A. Sharma, D.A. Karagodin, J. Jenrette, S. Gamez, P. Miesen, A. Caccone, M.V. Sharakhova, Z. Tu, P.A. Papathanos, R.P. Van Rij, O. S. Akbari, J. Powell, A. M. Phillippy, Bonizzoni M.

## Abstract

The Asian tiger mosquito *Aedes albopictus* is globally expanding and has become the main vector for human arboviruses in Europe. Here we present AalbF2, a dramatically improved assembly of the *Ae. albopictus* genome that has revealed widespread viral insertions, novel microRNAs and piRNA clusters, the sex determining locus, new immunity genes, and has enabled genome-wide studies of geographically diverse *Ae. albopictus* populations and analyses of the developmental and stage-dependent network of expression data. Additionally, we built the first physical map for this species with 75% of the assembled genome anchored to the chromosomes. These up-to-date resources of the genome provide a foundation to improve understanding of the adaptation potential and the epidemiological relevance of this species and foster the development of innovative control measures.

**One Sentence Summary:** Long-read and Hi-C-based *de novo* assembly of the arboviral vector *Aedes albopictus* genome fosters deeper understanding of its biological features.

## Main Text

Climate change, urbanization and increased international mobility are predicted to further increase the spreading of the highly invasive mosquito *Aedes albopictus* and severely exacerbate the risk and burden of *Aedes*-transmitted human pathogens, *in primis* dengue, Zika and chikungunya viruses, but also the veterinary-relevant parasite *Dirofilaria immitis* (*1*, *2*). As a consequence, nearly a billion people could face their first exposure to arboviral transmission within the next century especially in subtropical and temperate regions of the world, including Europe (*2*).

The initial genome assembly of *Ae. albopictus* (AaloF1) from the Chinese Foshan strain represented a fundamental achievement for the genetic characterization of this mosquito (*3*). From this analysis, based solely on the assembly of short DNA sequence reads, the genome of *Ae. albopictus* appears to be the largest mosquito genome sequenced to date (1.9 Gb). However, due to very high levels of repetitive DNA and reliance on short-read sequencing, AaloF1 remains highly fragmented with more than 150,000 scaffolds, limiting its utility.

Using a cytofluorimetric approach, we estimated the genome length of *Ae. albopictus* to be similar to that of *Ae. aegypti*, between 1.190-1.275 Gb, across populations from the native home range (Thailand, Malaysia, Singapore), old-colonized regions (La Reunion Island), and recently invaded areas (Italy, USA and Mexico) (Fig. 1A). To foster continuity, we chose to use the Foshan strain for further genome study. After six consecutive rounds of single sister-brother matings, we extracted high molecular weight DNA from forty sibling mosquitoes. We then generated approximately 82 Gb of PacBio single molecule long reads with a mean read length of 10 kb and an N50 length of 18 kb (N50 length: half of the data comprises sequences of this length or longer). Additionally, we prepared a Hi-C proximity ligation library from ten adult mosquitoes and collected 135 Gb of Illumina reads. We assembled the long-read PacBio data with Canu (*4*) and polished the resulting contigs with Arrow (https://www.pacb.com/products-and-services/analytical-software/smrt-analysis/) using the raw PacBio signal data. This initial assembly totaled 5.17 Gb, far exceeding the expected haploid genome size (~1.25 Gb), suggesting the presence of artificially duplicated alleles in the assembly. We hypothesized that this was due to high levels of heterozygosity in the pool of sequenced mosquitoes, resulting in multiple allelic variants assembled separately, as has been previously noted in long-read assemblies (*5*). To partition this initial assembly into primary and alternative contig sets, we analyzed contig alignments and depth of coverage with Purge Haplotigs (*5*) along with BUSCO single-copy orthologs (*6*) to determine which contigs were likely to be redundant and should be designated as alternative alleles. Haplotig purging reduced the size of the primary assembly by nearly half to 2.54 Gb, which was then scaffolded via the Hi-C data using SALSA2 (*7*).

**Fig. 1.**
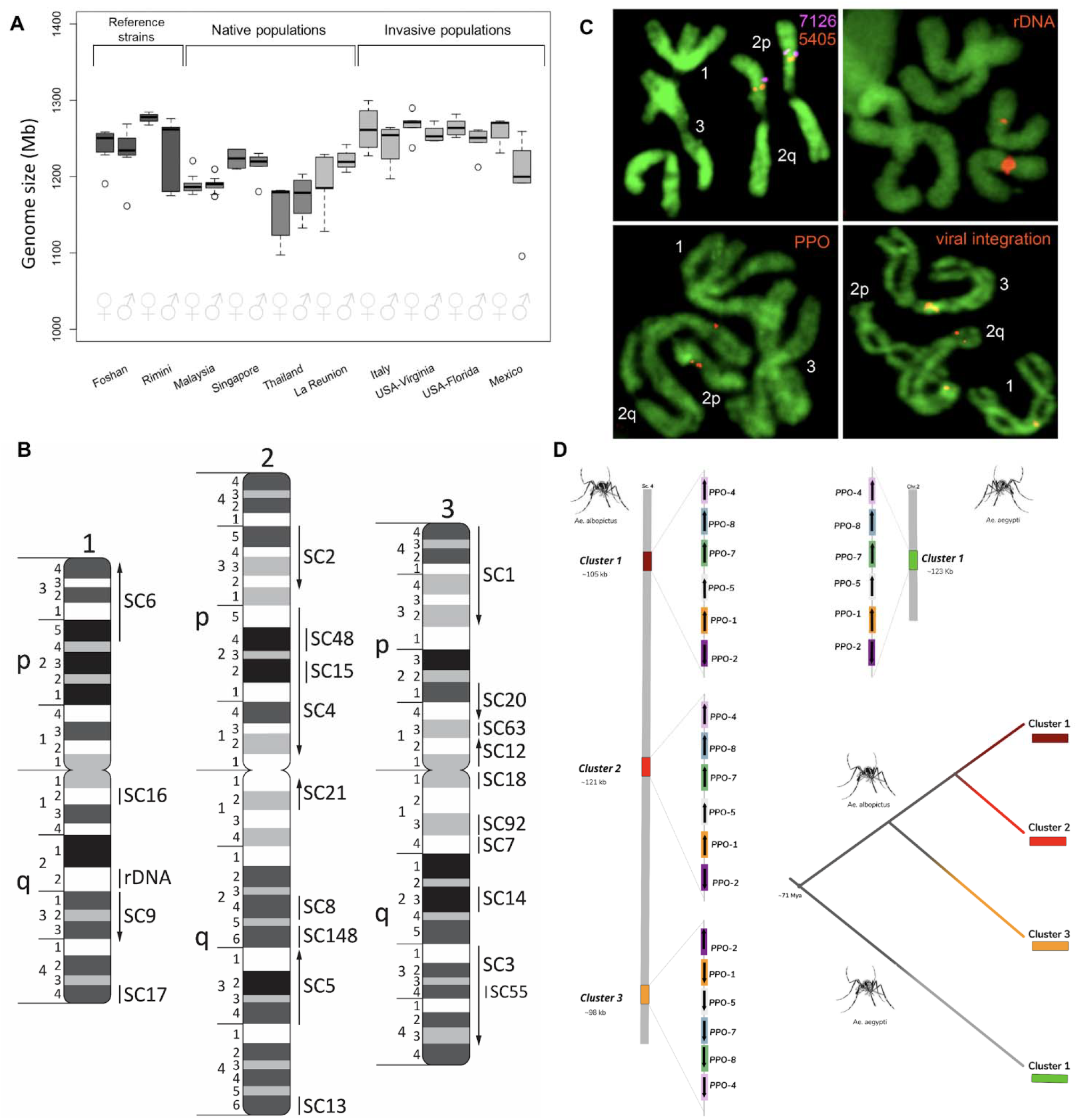
Size of the *Aedes albopictus* genome and physical map. **A)** Cytofluorimetric-based estimates of the genome size of *Ae. albopictus* strains, including Foshan and Rimini from which genome assemblies were derived based on short-read Illumina sequencing (*3*, *58*) and *Ae. albopictus* wild-collected samples from the native home range (Malaysia, Singapore, Thailand), an old-colonized region (La Reunion), and newly invaded areas (USA, Mexico, Italy). The *Ae. albopictus* genome size is estimated to be in the range of 1095-1299 Mb, comparable or slightly larger than that of *Ae. aegypti* (1066-1309 Mb) (*9*). **B)** Physical genome map of *Ae. albopictus* based on 50 DNA probes hybridized *in situ* to mitotic chromosomes. Chromosomes and chromosome arms are indicated by numbers 1, 2, 3, and letters p, q, respectively. Chromosome divisions and subdivisions are shown on the left sides of the ideograms. Scaffolds are indicated by arrows or lines. Arrows indicate orientations of the scaffolds. SC stands for scaffold; rDNA stands for ribosomal locus. **C).** Examples of fluorescence *in situ* hybridization. Chromosomal locations of transcripts XM_019675405 and XM_020077126 from scaffold 4 and 48, respectively; rDNA; polyphenol oxidase (PPO) gene clusters; and the largest viral integration in the genome (Canu-Flavi19) are demonstrated. Transcripts are indicated on a figure by the last four digits of their accession numbers. **D)** Schematic illustration of chromosomal locations of PPO cluster triplication in the new assembly of *Ae. albopictus* (GCF_006496715.1). Comparative genomics analysis of the synteny in AaegL5 genome reveals an array of 6 genes localized in a region of 123.44 Kb at chromosome 2 (2:199,230,485-199,353,929) was locally duplicated twice in *Ae. albopictus*, resulting in 18 PPO genes. PPO gene cluster array of chromosome 2 of *Ae. aegypti* includes AAEL015116 (PPO1), AAEL015113 (PPO2), AAEL013492 (PPO5), AAEL013493 (PPO7), AAEL013501 (PPO4), AAEL013496 (PP08).

The final primary assembly, which we call AalbF2, consists of 2,197 scaffolds with an N50 length of 55.7 Mb (Table S1). This represents a continuity increase of two orders of magnitude compared to AaloF1 scaffold N50s of 201 kb (*3*). This significant increase in continuity provides a more complete view of the genomic organization of *Ae. albopictus* and allows for a more accurate annotation of gene structures.

Analyses of single-copy orthologues via BUSCO (*6*) in AalbF2 showed an 8.3% increase in the percentage of complete, single-copy BUSCOs with respect to AaloF1 (Table S1). AalbF2 has a BUSCO completeness of 93.2%, with an estimated 14.0% duplication. Additionally, using Barrnap (https://github.com/tseemann/barrnap), the number of ribosomal RNA gene sequences was estimated to be 484 in AalbF2 (compared to 22 in AaloF1), a value close to the number (430) independently estimated from an *Ae. albopictus* haploid genome (*8*). The rate of alignments of DNA and RNA sequencing data from published resources, and the percentage of properly paired reads, were also analyzed and confirmed the quality and continuity of AalbF2 (Table S1). The higher continuity of AalbF2 is also shown by the annotation of transposable elements (TE), which amount to 55.03% of the genome size, a value comparable to that of the most recent assembly of the *Ae. aegypti* genome, AaegL5 (Table S2).

The original, unfiltered and unsplit assembly (main and alternative scaffolds) had a BUSCO completeness of 97.6% with 81.8% duplication, indicating that the majority of genes were represented in the combined assembly by more than one allele. Consistent with other long-read assemblies of other *Aedes* spp. mosquitoes, the large size of the genome coupled with small quantities of high molecular weight DNA obtainable from a single individual requires pooling of heterozygous individuals and the necessity of removing haplotypic duplications prior to the creation of haploid reference scaffolds (*9*).

A total of 26,856 protein-coding sequences were predicted in AalbF2 through the NCBI Eukaryotic Genome Annotation Pipeline (https://www.ncbi.nlm.nih.gov/genome/annotation__euk/process/). To help distinguish between artifacts and genuine gene duplications, which are resistant to proper assembly, and mitigating the heterozygosity effect from the original pooled DNA, we developed a pipeline based on the assumptions that selection acts mainly on the coding sequence of a gene and that homology between highly related paralogs drops in the flanking untranslated sequences (Fig. S1). To perform the analysis, we compared 500 bp or 1,000 bp of the flanking regions at the 5’ and 3’ ends of all candidate gene duplicate with an all-against-all BLASTn search with an e-value of 1×10^−40^ for each flanking region. We found 1,329 (8.05% of total) genes with high similarity within 500 bp of their 5’ and 3’ flanking regions mapping on 452 of the 2,196 scaffolds (Supplementary Material). When we considered the extended 1,000 bp regions, the number of candidates duplicated was lower (808 mapping on 300 scaffolds; 4.89 % of total). Most of these artifacts involved a single duplicated gene (twins) and the number decreased with increasing copies. A list of gene duplications that are likely to be artifacts of the assembly is available for future reference in Database S1.

A significant improvement of AalbF2 is that more than 50% of the genome assembly is contained within the 13 largest scaffolds (e.g. L50=13; L75=58, Fig. S2, Table S1). We developed a physical genome map of the *Ae. albopictus* genome using *in situ* hybridization on mitotic chromosomes covering 57% of the genome assembly by targeting twenty of the largest genomic scaffolds and three minor scaffolds (Table S3). We developed a new mapping approach based on amplification of DNA probes using cDNA instead of Bacterial Artificial Chromosome (BAC) clones and hybridized fifty transcript fragments or gene exons from twenty-four scaffolds to mitotic chromosomes, thus generating the first physical map of the *Ae. albopictus* genome (Fig. 1C). Target probes were derived from transcripts of the *Ae. albopictus* C6/36 cell line genome (*10*). For most scaffolds, at least a pair of DNA probes was amplified, labeled with different dyes and hybridized simultaneously to the chromosomes. Use of alternative dyes allows not just mapping, but also orienting of the genomic scaffolds along the chromosomes. All DNA probe pairs hybridized to the same chromosomes at the distance proportional to the size of the genomic scaffolds indicating that the genomic scaffolds were assembled correctly. The positions of the probes were assigned to the chromosome bands on ideograms (Fig 1C). A total of 4, 9, and 10 scaffolds were assigned to the chromosome 1, 2 and 3, respectively. Positions of the transcript from scaffolds 15, 48, and 55 hybridized to places already covered with other large scaffolds. The positions of all tested transcripts were consistent with their positions in the *Ae. aegypti* genome, which is assembled into chromosome-size scaffolds, providing an independent confirmation of the accuracy of the *in situ* hybridization results (*9*). Based on probe-mapping to the *Ae. aegypti* genome and homology between the *Ae. aegypti* and the *Ae. albopictus* chromosomes (Fig. S2), we bioinformatically assigned the 58 longest scaffolds covering 75% of the genome to *Ae. albopictus* chromosomes (Table S4).

Cytogenetic comparison (Table 1) between *Ae. albopictus* and *Ae. aegypti* demonstrated that the total chromosome length is 4.9 µm or 16.4% longer in *Ae. albopictus* (P<0.0001), which suggests a slightly larger genome size in this species, as also suggested by cytofluorimetry. Chromosome proportions, such as relative chromosome and arm lengths, between the two species were also different. In *Ae. albopictus* “chromosome 1” was shorter but chromosome 2 was longer relative to *Ae. aegypti*. Besides positioning and orienting the largest scaffolds, we physically mapped the18S rDNA and other genomic features (e.g., viral integrations and representative immunity genes) described below (Fig. 1C, D). The 18S rDNA mapped in the region of the secondary constriction in region 1q22. The intensity of the signal significantly varied among chromosomes from individual mosquitoes suggesting variations in numbers of ribosomal genes.

**Table 1.**
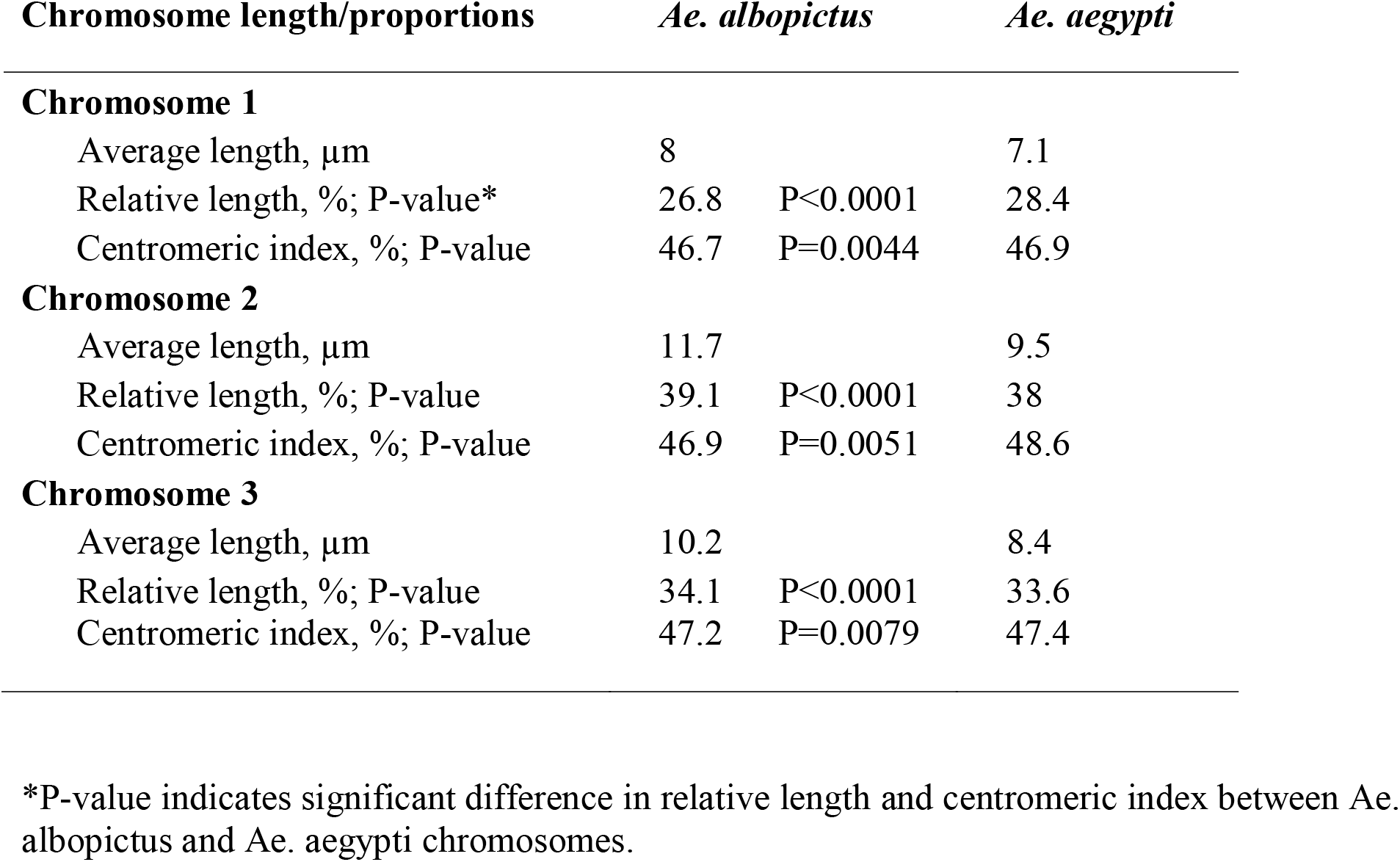
Comparison between *Ae. aegypti* and *Ae. albopictus* mitotic chromosomes.

| Chromosome length/proportions | *Ae. albopictus* |  | *Ae. aegypti* |
| --- | --- | --- | --- |
| **Chromosome 1** |  |  |  |
| Average length, μm | 8 |  | 7.1 |
| Relative length, %; P-value* | 26.8 | P<0.0001 | 28.4 |
| Centromeric index, %; P-value | 46.7 | P=0.0044 | 46.9 |
| **Chromosome 2** |  |  |  |
| Average length, μm | 11.7 |  | 9.5 |
| Relative length, %; P-value | 39.1 | P<0.0001 | 38 |
| Centromeric index, %; P-value | 46.9 | P=0.0051 | 48.6 |
| **Chromosome 3** |  |  |  |
| Average length, μm | 10.2 |  | 8.4 |
| Relative length, %; P-value | 34.1 | P<0.0001 | 33.6 |
| Centromeric index, %; P-value | 47.2 | P=0.0079 | 47.4 |

\*P-value indicates significant difference in relative length and centromeric index between *Ae. albopictus* and *Ae. aegypti* chromosomes.

### The landscape of endogenous viral elements

Among all Culicinae mosquitoes, the genomes of *Ae. aegypti* and *Ae. albopictus* have an exceptionally high count of nonretroviral RNA virus sequences that are integrated into repetitive genomic regions. Both genomes contain several hundred of these so called nonretroviral Endogenous Viral Elements (nrEVE) or Nonretroviral Integrated RNA Virus Sequences (NIRVS) (*11*). The contiguity of AalbF2 enables an improved understanding of the landscape of viral integrations in the *Ae. albopictus* genome. Using a viral database composed of 1,563 viral species (Database S2), a total of 456 viral integrations were identified in AalbF2 and classified into ten taxonomic categories (Fig.2A, Database S3). Correspondence between viral integrations previously annotated in AaloF1 was evaluated (*11*), considering that the larger database used for AalbF2 led to a higher number of nrEVEs (Table S5). Additionally, we used the identified viral integrations to screen the alternative assembly (NCBI accession GCA_006496715.1) and found alternative nrEVE alleles (Table S6), confirming that the haplotig purging applied to the initial assembly effectively moved haplotypic variants into the alternative assembly.

The largest viral integration (Canu-Flavi19) in AalbF2 reached 6,593 bp and encompassed all of the structural proteins, the entire NS1 and NS2 and part of the NS3 and NS5 proteins of the 11,064 bp genome of Aedes Flavivirus, with a 97,63 percentage of identity (Database S3). This viral integration was mapped by *in situ* hybridization to chromosome 2q close to the telomere, confirming it is integrated within the genome (Fig. 1B). Signals were also found in the centromeres of all three chromosomes, probably because these regions contain nrEVEs with sequence similarity to Canu-Flavi19 (Fig. 1B). In total, viral integrations occupied 585,891 bp (0.02% of the total genome), spread across 115 out of the 2,197 assembled scaffolds and were distributed homogeneously across scaffolds. A total of 27 and 67 viral integrations were identified with similarities to Flaviviruses and Rhabdoviruses, respectively (Database S3). All Flavi-EVEs have similarities to known insect specific viruses (ISVs), primarily Aedes Flavivirus (AeFV), La Tina Virus (LTNV), Cell Fusing Agent Virus (CFAV) and Kamiti River Virus (KRV). These ISVs are at the base of the phylogenetic tree of Flaviviruses (*12*). When all Flavi-EVEs are mapped against a representative Flavivirus genome, they cover the whole genome, with overrepresentation of the E and NS5 coding regions (Fig. S3). Overrepresentation is due both to duplications and independent integrations as shown by the high range of identity to the E and NS5 proteins of circulating viruses, accompanied by similarities to different viral species (Database S3). Most Rhabdo-EVEs are similar to the glycoprotein coding region of different Rhabdoviruses (Fig.S3). Both Flavi- and Rhabdo-EVEs tend to map less than 10kb to each other, generating clusters. Clusters may include both Flavi- and Rhabdo-EVEs or be composed uniquely of viral integrations from one viral family (Fg. S3). Viral integrations are tightly associated with transposable elements (TE), primarily Gypsy and Pao LTR retrotransposons, as seen in AaloF1 (Fig. 2C). This association appears to be driven by the enrichment of LTR retrotransposons into piRNA clusters (Fig. S3).

**Fig. 2.**
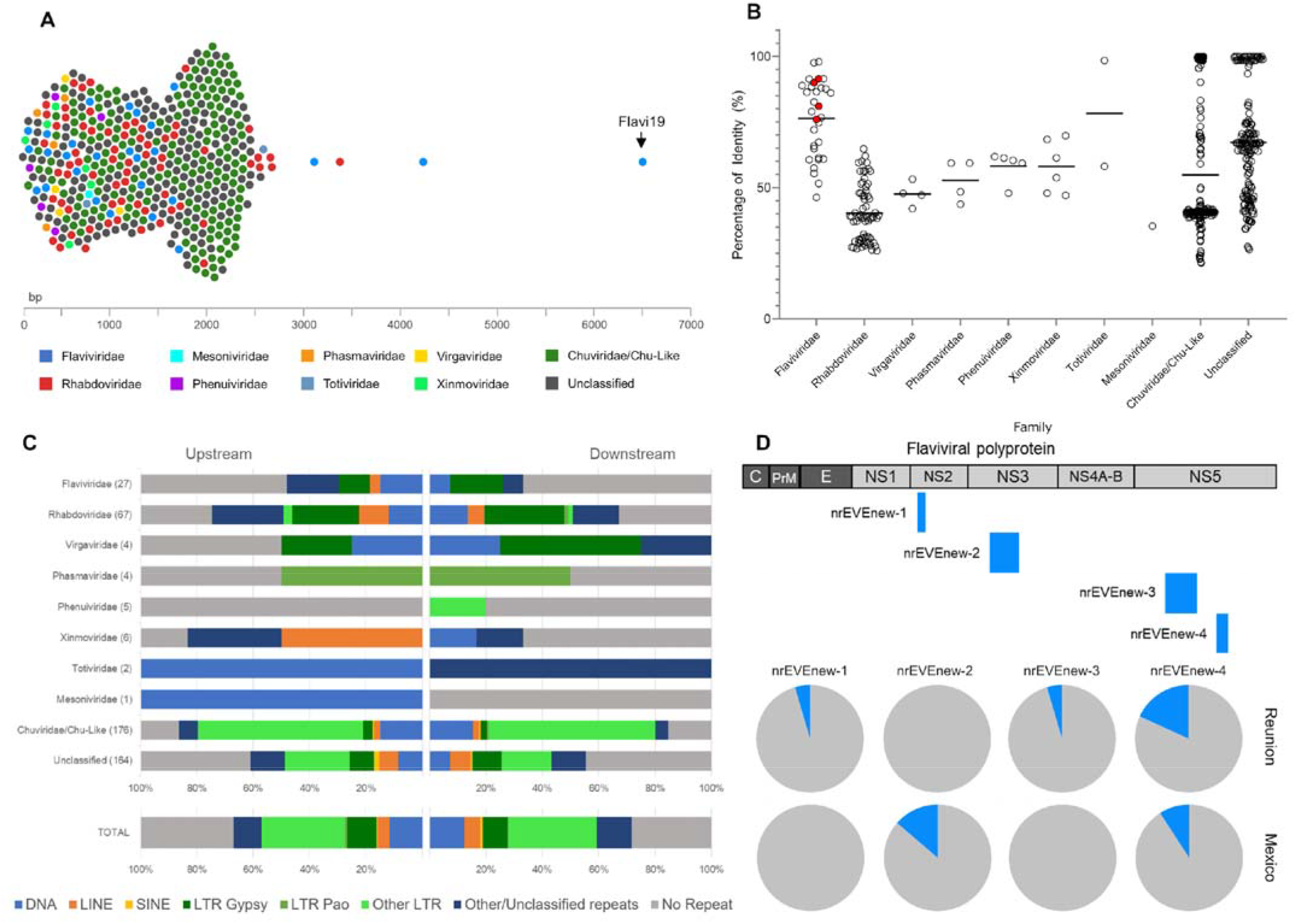
Atlas of viral integration in the *Aedes albopictus* genome. **A)** Beeswarm plot showing viral integrations identified in the *Ae. albopictus* genome (AalbF2). Each dot is a viral integration, plotted according to its length and color-coded based on its viral origin. nrEVEs range in length from 131 to 6593 nt, with an average of 1289 nt. Arrow points to Canu-Flavi19, the longest nrEVE. **B)** Scatter plot representing the amino acid identity of each nrEVE and its best hit retrieved by blastx searches against NR database grouped by viral family. The average is shown by a line. Red dots are the novel viral integrations discovered in wild-caught mosquitoes. **C)** Bar plots showing the type of closest transposable element, which was identified upstream and downstream each nrEVE. Viral integrations are classified based on their viral origin as shown in panel A. **D)** Scheme of the novel viral integrations identified in the genome of wild-collected mosquitoes with respect to a Flavivirus genome and their frequency occurrence in mosquitoes from Tampon (Reunion) and Tapachula (Mexico).

Viral integrations have been proposed to be adaptive heritable immunity sequences, which implies that their distribution patterns may differ across geographic populations depending on viral exposure (*11*, *13*, *14*). To address whether viral integrations different from those annotated in AalbF2 can be characterized in wild-caught mosquitoes (hereafter called novel nrEVEs), we collected and sequenced the genomes of 24 adult females from Tapachula (Mexico) and Tampon (La Reunion island) where several arboviruses are endemic. By using Vy-Per (*15*) and custom scripts, we identified one and two novel viral integrations in samples from Tapachula and Tampon respectively, plus a novel viral integration common to both populations (Fig.2D). Two of these novel viral integrations (nrEVEnew-3 and nrEVEnew-4) have similarities to AeFV, one (nrEVEnew-2) to CFAV and the other (nrEVENew-1) to KRV (Fig.2D). All these novel viral integrations were molecularly validated by designing specific PCR primers (TableS7, Fig. S3B). Novel viral integrations were more frequent in mosquitoes from Tampon than Tapachula (Fig.2D). Additionally, two of the Tampon novel viral integrations had a 90% amino acid identity with AeFV and CFAV, respectively (compared to the 72% average identity for annotated Flavi-EVEs), suggesting recent integration events. This result correlates with the invasion history of *Ae. albopictus* out of its native home range in Asia. Before the aggressive global invasion of *Ae. albopictus*, which started roughly 50 years ago, *Ae. albopictus* had reached the islands of the Indian and Pacific Ocean from South East Asia in the 18^th^ to early 20^th^ centuries (*16*). Thus, mosquitoes from La Reunion island are considered “old” and have maintained large populations (*17*, *18*). In contrast, *Ae. albopictus* was first detected in Tapachula in 2002, likely a secondary invasion from USA or Italy (*19*, *20*).

### Distribution and structure of piRNA clusters

PIWI-interacting RNAs (piRNAs) are mostly known for their role in immunity against TEs in the germline (*21*). This is best studied in the model organism *Drosophila melanogaster*. However, in *Aedes* spp. mosquitoes, the piRNA pathway acquired additional functions in antiviral immunity, and can use viral RNAs as substrate for piRNA production (*22*). Most piRNAs are derived from large genomic regions termed piRNA clusters. These clusters present a memory of past transposon invasions and confer immunity against these elements, as piRNAs processed from transposon remnants within clusters can target active transposons encoded elsewhere in the genome (*21*).

Using the preceding AaloF1 genome assembly, a previous study reported 643 clusters with a maximum length of maximum 10 kb (*23*). However, piRNA clusters can span up to several hundred kilobases (*24*, *25*), therefore, a more continuous genome assembly can improve annotation of these genomic regions. We used small RNA libraries generated from somatic tissues (female carcass) as well as germline tissues (ovaries) (Supplementary Material) to annotate 1441 piRNA clusters with an average size of 10.911 kb (SD 634.885 kb; max: 139.92 kb) (Fig. S4, Database S4), covering 0.62% of the genome. This is comparable to piRNA clusters annotated with the same approach in *Ae. aegypti* (Fig. 3A). In contrast, using the same annotation pipeline on the highly fragmented *Ae. albopictus* AaloF1 genome assembly, we recovered nearly twice as many (2467) but much smaller clusters (average size: 5.923 kb; SD: 306.239 kb; max 64.225 kb) (Fig. 3A, B). Only a comparably small fraction (31.8% and 47.3%) of all piRNAs in the germline and soma, respectively, were included in piRNA clusters in AalbF2, while this fraction was nearly twice as large in *Ae. aegypti* (Fig. 3A). This is likely accounted for by the 14% of duplications still present in the assembly, leading to the exclusion of piRNA clusters without, or with only very few uniquely mapping piRNAs, the presence of which was used as a criterium to annotate piRNA clusters. Consequently, when only considering unambiguously mapping piRNAs, the fraction of piRNAs included in clusters increases to 59.1 % and 72.9 % in germline and soma, respectively.

**Fig. 3.**
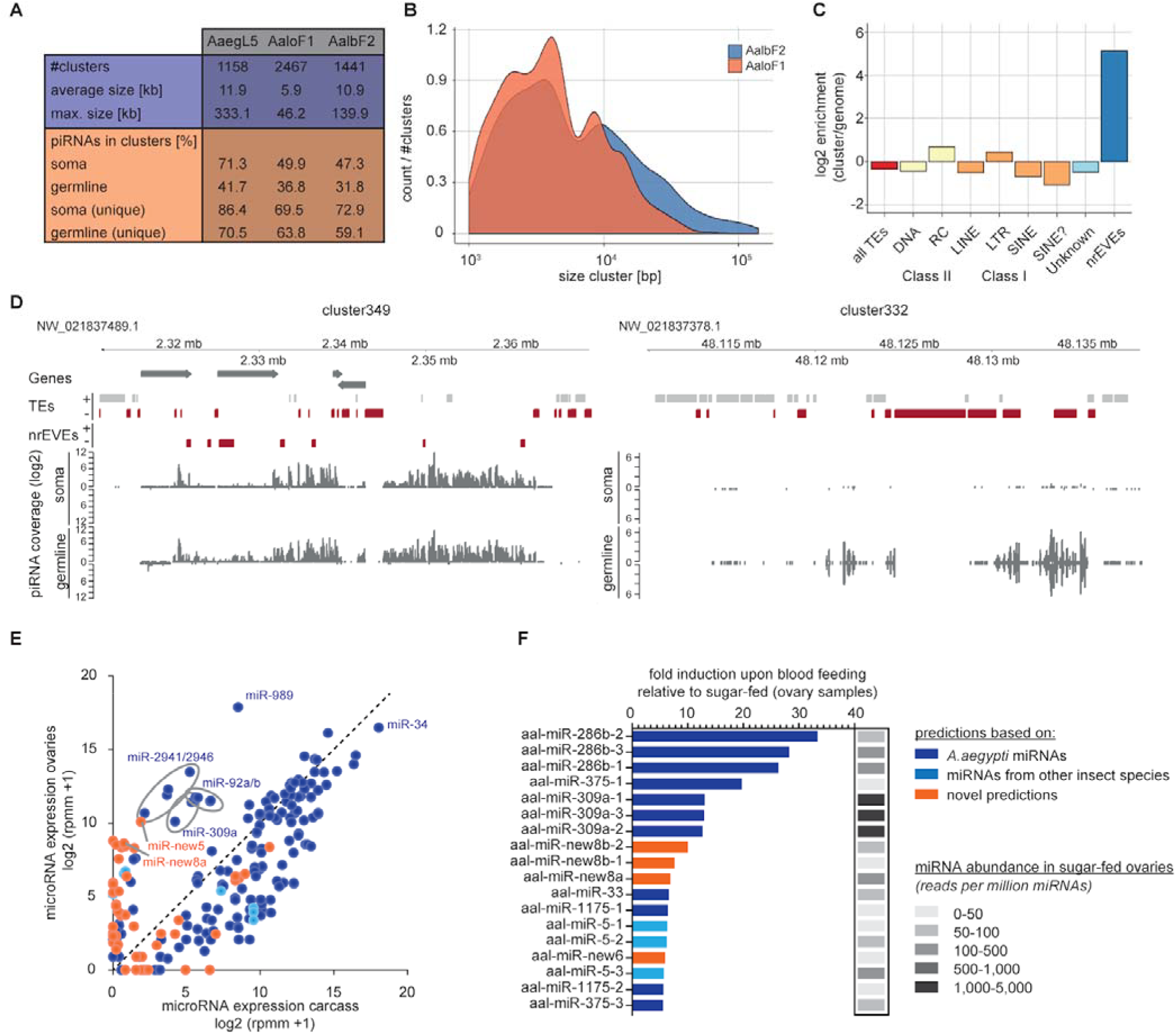
Small non-coding RNA annotation in AalbF2. **A)** Summary statistics on annotated piRNA clusters using the genome assemblies for *Ae. aegypti* AaegL5 and *Ae. albopictus* AaloF1 or AalbF2. **B)** Size distribution of piRNA clusters annotated with the old AaloF1 assembly or the most recent AalbF2 assembly. The density plot shows the number of clusters normalized to the total number of piRNA clusters. **C)** Enrichment of repeat classes or nrEVEs in piRNA clusters compared to the whole genome. **D)** log2 piRNA coverage on an exemplary uni-strand (left panel), or dual-strand (right panel) piRNA cluster (given as piRNAs per million mapped reads [rpm]). Annotated genes are indicated with arrows, repeat features and nrEVEs with gray or red boxes for positive or negative strands, respectively. **E)** miRNA abundance in *Ae. albopictus* carcass and ovary samples. Counts for individual miRNAs were normalized to the total number of miRNAs in each dataset and expressed as log2 transformed reads per million miRNAs (rpmm) + 1. The mean of two independent libraries for each condition is shown. The highly abundant miR-34 and selected miRNAs with high expression in ovary, but not carcass are indicated. **F)** Fold induction of miRNA levels in blood-fed ovary samples compared to sugar-fed samples. Basal expression of each miRNA in the sugar-fed samples is indicated in grey scale. Only miRNAs with an induction ≥ 5-fold are shown. Color coding in A and B represent the basis for the miRNA prediction, as indicated.)

The vast majority of all clusters display piRNA expression biased towards one strand, and only approximately one fifth of all clusters were expressed from both strands (see exemplary clusters in Fig. 3C). Such dual strand clusters were mostly expressed in the germline (Fig. S4). Interestingly, relative piRNA expression from clusters varied substantially between somatic and germline tissues, with some clusters showing a soma-dominant expression and others being predominantly expressed in the germline. Blood-feeding had little impact on cluster expression. Analysis of publicly available small RNA libraries derived from the widely used *Ae. albopictus* C6/36 and U4.4 cell lines showed piRNA production from both somatic and germline clusters (Fig. S4).

While piRNA clusters are highly enriched with transposable elements in fruit flies (*25*), this is not the case in *Ae. aegypti* mosquitoes (*26*), even though their genomic transposon content is much higher. Comparably, only a minority of *Ae. albopictus* piRNAs were derived from repetitive elements (*23*), and piRNA clusters were slightly depleted of all repetitive sequences except for helitrons and LTR-retrotransposons (Fig. 3C). Interestingly, nrEVEs were enriched compared to the rest of the genome (Fig. S3), and 138 out of 456 elements were overlapping with piRNA clusters, suggesting strong evolutionary pressure to integrate viral sequences into piRNA clusters and/or maintain nrEVEs in piRNA-producing loci.

### miRNA annotation

Small-noncoding RNA pathways contribute to important biological and cellular processes like development, differentiation, and immunity. MicroRNAs (miRNAs) are an endogenous class of small regulatory RNAs that are crucial for post-transcriptional regulation of gene expression (*27*). MiRNAs are processed from precursor hairpin structures (pre-miRNAs) which are present in the genome as single copy loci or, due to gene duplication, as multiple copies of the same miRNA. A comprehensive inventory of *Ae. albopictus* miRNAs is an important resource for investigating small RNA function in vector biology and mosquito antiviral immunity. The official depository of miRNA genes across all species, miRbase (*28*), does not currently include *Ae. albopictus* miRNAs. Therefore, to annotate miRNA genes in AalbF2, we used the miRDeep2 algorithm (*29*) on data from small RNA libraries as described above, comprising of more than 23 million miRNA-sized 18-24nt reads (Supplementary Material). The majority of reads were derived from carcass samples, which is expected as small RNA libraries prepared from ovary samples are more biased towards piRNAs. Initially, miRDeep2 predicted 473 pre-miRNA loci in AalbF2, which was reduced to 229 loci representing 121 distinct pre-miRNA species (Database S5) after manual inspection and handling stringent prediction criteria (Supplementary Material). Amongst these predictions, 92 represent miRNAs previously annotated in the *Ae. aegypti* genome, three were predicted based on conservation to miRNAs in other insect species, and 26 were entirely novel miRNA genes. Using these predictions, we characterized the expression of miRNAs in ovaries and carcasses and analyzed changes induced by blood-feeding. We found that most highly abundant miRNAs show a similar expression pattern between ovaries and carcass (Fig. 3E). Yet, a group of miRNAs, including miR-92a/b, miR-309a, miR-989, miR-2941, miR-2946 and a newly predicted miRNA, miR-new5, were highly abundant (>1000 reads per million miRNAs; rpmm) exclusively in the ovary samples (Fig. 3E). These findings are coherent with previous studies that identified the clustered miRNAs miR2941/2946 to be specifically expressed in *Ae. aegypti* ovaries (*30*). miR-989 is known to be amongst the most abundant miRNA in mosquito ovaries, both in *Anopheline* and *Aedes* spp. mosquitoes (*31*, *32*). Similarly, miR-309 was found to be predominantly expressed in *Ae. aegypti* ovary tissue and was furthermore shown to be strongly induced upon blood feeding both in *Aedes* and *Anopheles spp.* mosquitoes (*33*, *34*). When comparing sugar and blood-fed *Ae. albopictus*, we observe a similar induction of miR-309a upon blood feeding (Fig. 3F). Likewise, miR-286b and miR-375, which we find to be strongly induced upon blood meal, have previously been shown to be upregulated after blood meal in *Anopheles stephensi* and *Ae. aegypti*, respectively (*34*, *35*), indicating that an orchestrated miRNA response to blood feeding is conserved between different mosquito species. We noted that most newly predicted miRNAs are predominantly expressed in ovary tissue (Fig. 3E), which likely reflects a sampling bias of previous studies that did not deep-sequence and predict miRNAs from dissected ovary samples. Some of these predicted miRNA species are relatively highly abundant and are differentially expressed upon blood feeding, suggesting important function in the physiological processes that are induced upon blood meal.

### Curation of immunity repertoire

The capacity of mosquitoes to acquire, disseminate and transmit viruses (i.e. vector competence) is a complex phenotype which is controlled by genetic elements of both the vector and the pathogen, as well as environmental variables (*36*). Understanding the complex relationship between vectors and pathogens requires understanding innate immunity in mosquitoes. To catalog genes encoding the immune repertoire of *Ae. albopictus* we searched with BLASTp the predicted peptides of the AalbF2 assembly using as a query 417 manually curated proteins of *Ae. aegypti* from ImmunoDB (*37*). We combined phylogenetic comparisons and manual annotation to curate 663 putative immune-related genes encoding 979 predicted proteins, belonging to 27 functional groups (Table 2A, Database 6). This value is in line with that estimated in AaloF1 (521 genes), confirming the finding that the immune repertoire of *Ae. albopictus* is larger than that of other dipteran species (*3*, *37*). A manual inspection of the 663 putative immune-related genes using our 5’ and 3’ flanking region pipeline identified a set of 78 suspicious genes that are distributed in half of the immune gene families (Table 2 and Database 6), reducing the total number of predicted immune genes to 622.

**Table 2.**
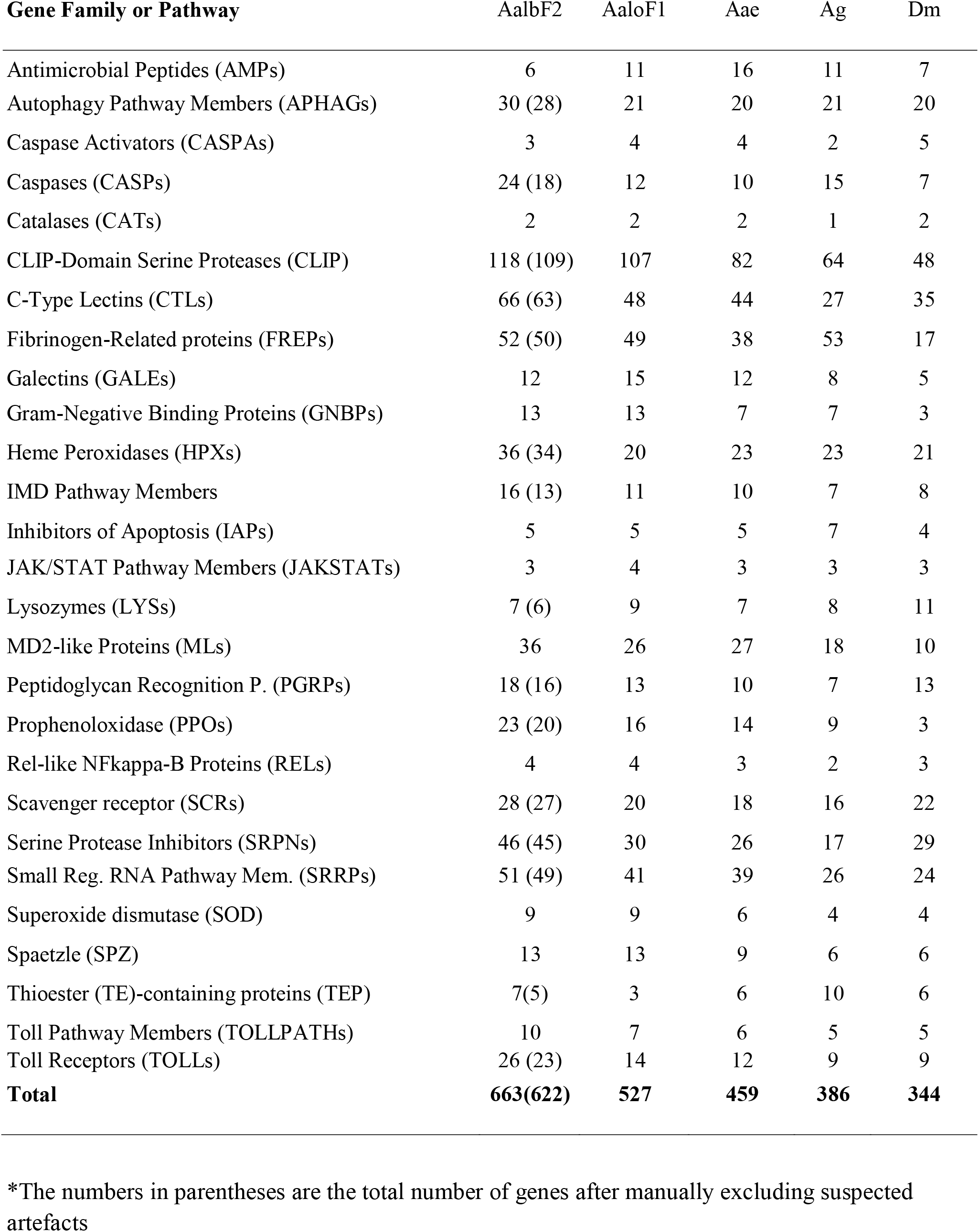
The repertoire of immune genes of *Aedes albopictus*. Comparison in the number of immune-related genes among *Aedes albopictus* (Alb, AalbF2 and AaloF1 assemblies), *Aedes aegypti* (Aae, AaegL5), *Anopheles gambiae* (Ag, AgamP4) and *Drosophila melanogaster* (Dm, Dmel_r6.29).

Immune system functions can be broadly categorized into three main phases, recognition, signal transduction and effectors (*37*–*39*). A detailed analysis of the immune repertoire of *Ae. albopictus* revealed extensive expansions in 16 of the 27 functional groups relative to *Ae. aegypti.* In the Toll and IMD pathways, genes involved in recognition and Toll-1/Spz signal transduction show expansion, whereas immune effectors do not display similar family-wide augmentations. Interestingly, while five cecropins (CEC) genes are known in *Ae. aegypti*, we only identified a single CEC gene in the new assembly. We found expansions in families involved in all immune phases of the melanization pathway (*40*). The most extreme expansion event regards the CLIP family of regulators with 118 members compared to 67 and 56 genes reported for *Ae. aegypti* and *An. gambiae*, respectively (Table 2). Another interesting case involves the prophenoloxidase (PPO) gene family, which in *Ae. aegypti* includes six tandemly arrayed genes, namely PPO4, PPO8, PPO7, PPO5, PPO1, and PPO2. We found that the entire cluster of six genes has been locally duplicated twice in *Ae. albopictus*, resulting in 18 genes (Fig.1D, Table S8). We confirmed this triplication of the clusters using *in situ* hybridization (Fig.1B). PPOs are enzymes that catalyze the production of melanin in response to infection (*41*). Expansion of PPO genes is not common in insects (*42*), but in mosquitoes the number of genes is higher than other insects. The high conservation of the PPO-organization and order in the array in both *Ae. aegypti* and *Ae. albopictus* strongly suggests that these duplications are ancient events that occurred 71.4 Mya before the split between the two species (*3*). Future studies focusing on dissecting the functional importance of specific family expansions in *Ae. albopictus* may determine their significance for its biology including vector competence and ecological adaptation.

### The sex-determining M locus

In both *Ae. aegypti* and *Ae. albopictus*, sex is determined by a male-determining locus (M locus) that resides on one homolog of chromosome1. *Nix*, the dominant male-determining factor, was first discovered in the M locus of *Ae. aegypti* (*43*). We searched AalbF2 for *nix* and located it in an approximately 917 kb scaffold (NW_021838423.1). The *nix* sequence is male-specific as indicated by the Chromosome Quotient analysis (*44*) using Illumina reads obtained from male and female mosquitoes of the Foshan strain (*45*). A part of the *nix* gene was previously identified in *Ae. albopictus* (*43*, *46*) and its full-length sequence was described in the assembly of the *Ae. albopictus* C6/36 cell line (*10*). The *nix* gene in the AalbF2 assembly is annotated as having two exons flanking a small intron (XM_019669557.1), similar to a previous report (*9*). However, there is an apparently defective copy of *nix* approximately 22 kb away from XM_019669557.1. This copy does not have an intact open reading frame and fragments showed up to 70% amino acid identity to XM_019669557.1 (Fig. S5). Such a duplication has not been reported in *Ae. aegypti* (*43*). A second gene encoding a myosin heavy chain protein named *myo-sex* (*47*) has also been shown to be located in the M locus, together with *nix* in *Ae. Aegypti* (*9*). Myo-sex is required for male flight in *Ae. aegypti* (*48*). A *myo-sex* homolog (XM_019707039.1 or XP_019562584.1; Fig. S5) has been found in two separate contigs (NW_021838603.1 and NW_021838542.1). It is not yet clear whether the gene that encodes XP_019562584.1 is also located in the M locus in *Ae. albopictus*, as the Chromosome Quotient analysis (*44*) was complicated by the presence of highly similar autosomal paralogs (e.g., AALF000603 and XP_019560880).

### Genome-wide polymorphism and linkage disequilibrium

The level of genetic variability among populations of a given species is the substrate for evolution, which, for an invasive vector species like *Ae. albopictus*, includes processes of adaptation to new ecological settings, selection of resistance alleles against control tools (i.e. insecticides) and co-evolution with pathogens (*49*–*51*). These are biological features important to estimate the epidemiological relevance of *Ae. albopictus* populations and to account for in the design of novel genetic-based strategies of vector control (*36*, *52*). As for the analyses of the landscape of viral integrations, we used whole-genome sequencing (WGS) data of mosquitoes from Tapachula and Tampon (*53*) to show the usefulness of AalbF2 in understanding the genomic diversity of *Ae. albopictus* populations. The genetic diversity (π) estimates for the laboratory strain are lower than those for the wild populations, which is consistent with the hypothesis of a population bottleneck in the laboratory strain (Fig. 4A). Genetic diversity is slightly higher for the invasive Mexican population than the old population from La Reunion. Global estimates of genetic differentiation (*F*_*ST*_) among the three samples range from 0.13 to 0.21, with Foshan being the most differentiated (Fig. 4B). Sliding window analyses across the genome showed regions of high and low genetic differentiation between the two wild populations (Fig. 4C) and varying levels of genetic diversity for the two wild populations and the Foshan strain (Fig. 4D). We also derived estimates of linkage disequilibrium (LD). Across the three samples studied the *r^2^* Max/2 is approximately 1.3 kb (Fig. 4E). These estimates are strikingly smaller than estimated values for *Ae. aegypti*, which range between 34 to 101 (*9*). While comparing these LD estimates may be complicated by differences in data collection platforms (WGS for *Ae. albopictus* and SNP-chip for *Ae. aegypti*), the striking difference may reflect the different colonization histories of *Ae. aegypti* and *Ae. albopictus* populations (*16*, *54*). *Aedes aegypti* experienced a slow colonization process that started in the seventh century compared to a quick dispersal in the past 50 years for *Ae. albopictus* that resulted in genetic admixture among the invasive populations (*17*, *18*, *55*). The improved continuity of AalbF2 improves our ability to understand the spatial context of genetic signals and long-range patterns.

**Fig. 4.**
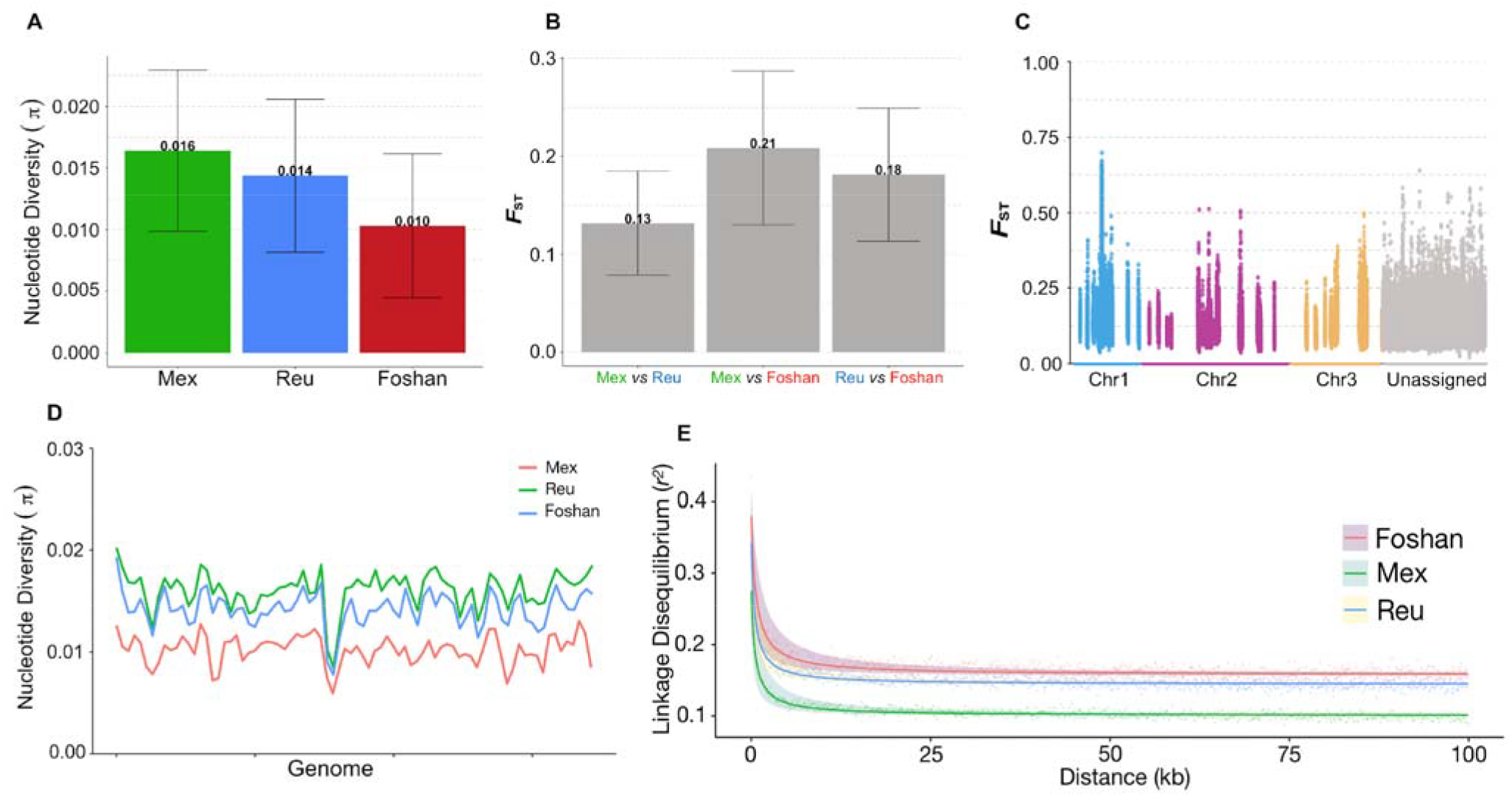
Genome-wide polymorphism in *Aedes albopictus.* **A)** Mean nucleotide diversity; **B)** Global *F*_*ST*_, the mean and standard deviation were calculated from sliding windows analysis. **C)** *F*_*ST*_ estimates between the two wild populations, Tapachula (green) and Tampon (Blue). *F*_*ST*_ estimates measured across the whole genome with sliding windows of 50kb with 10kb steps. Scaffolds that have been assigned to chromosomes 1, 2 and 3 are on the left side of the plot. The remaining unassigned scaffolds are shown on the right side. The unassigned scaffolds were placed in alphabetical order from left to right. The green line is the mean *F*_*ST*_ = 0.13 and the orange line is *F*_*ST*_ =0.46, which is 4 standard deviations above the mean. The highest peak near the center of the plot (red arrow) is found on scaffold NW_021838543.1. **D)** Overview of the pattern of nucleotide diversity across the genome. Nucleotide diversity was measured using 50Kb long sliding windows and 10kb steps. **E)** Linkage disequilibrium (*r^2^*) for two wild populations and a laboratory strain. Red, green and blue lines represent the fitting curve estimated with the *ngsLD* package and shaded areas around the lines represent confidence intervals from 100 bootstraps.

### Developmental Transcriptional Profile

Understanding the network of expression throughout development could provide insights into biological functions implicated in the adaptation of this invasive species to different environments and, coupled with the ability to manipulate genes and their expression, the basis to study gene function. Additionally, *cis*-regulatory elements that guide the expression in a tissue- or time-specific manner could be identified from the analyses of transcriptional profiles and be co-opted in novel genetic-based strategies of vector control. AalbF2 and its predicted gene models served the basis to establish a comprehensive global view of gene expression dynamics throughout *Ae. albopictus* development taking advantage of recently produced Illumina RNA sequencing (RNA-Seq) data from 47 unique samples representing 34 distinct stages of mosquito development (*56*) (Fig. 5A). These RNA-seq data amounted to 1.56 billion reads corresponding to total sequence output of 78.19 Gb (Database S7). A total of 94.1% of the reads were mapped to AalbF2. The number of spliced alignments increased substantially from 39,991,260 in the assembly of the C6/36 *Ae. albopictus* cell line (canu_80X_arrow2.2, *17*) to 56,243,825 in AalbF2 (40.64% increase), again confirming a more complete annotation in AalbF2 (Fig. 5B). The number of uniquely mapped reads also increased significantly likely due to the removal of extensively duplicated regions found in the C6/36 assembly (*10*).

**Fig. 5.**
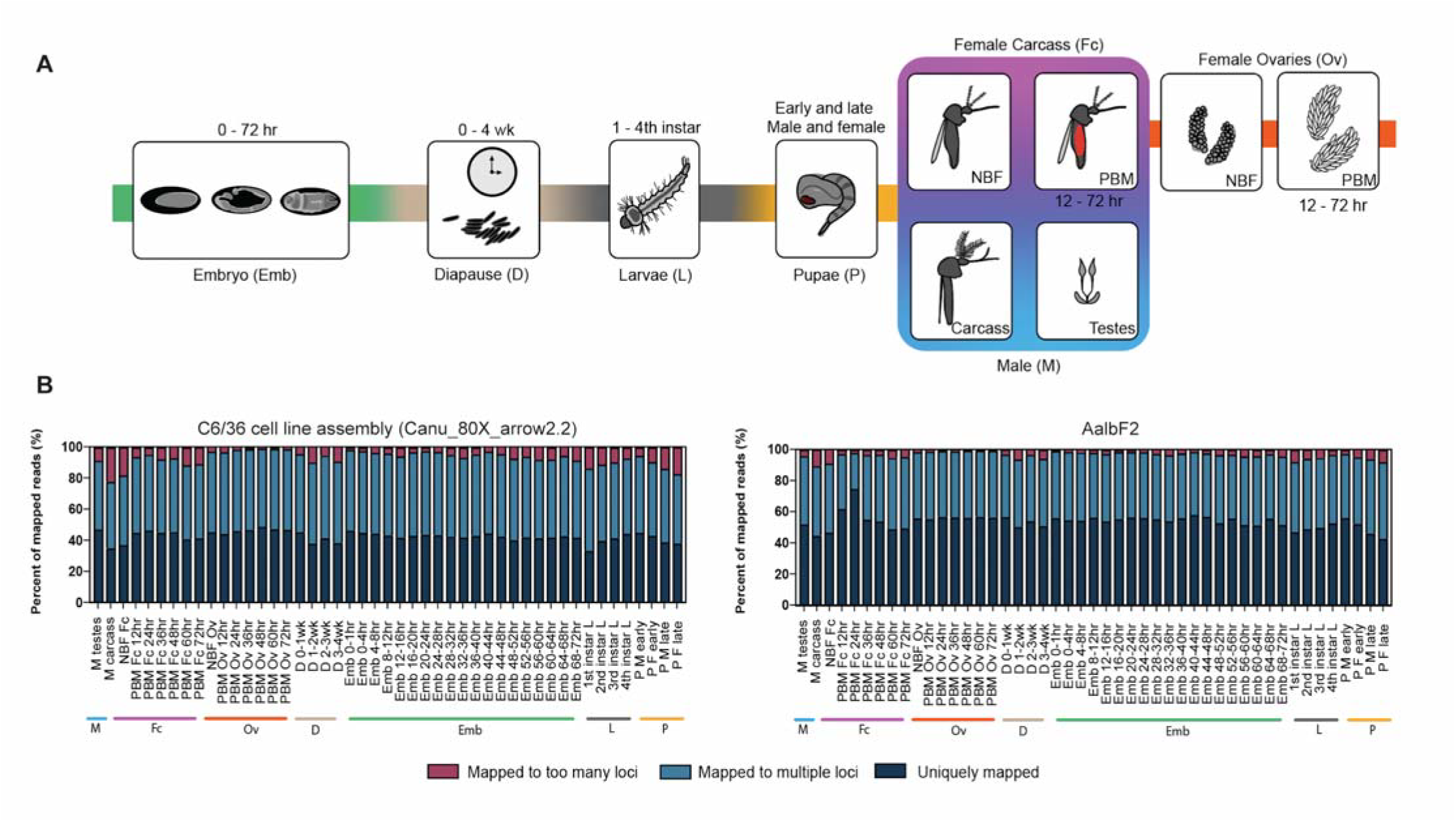
Schematic of mosquito development and mapping statistics of sequenced transcripts to *Ae. albopictus* genome assemblies. **A)** Developmental transcriptome of *Ae. albopictus*. Our developmental time-course included 34 stages spanning the major developmental groups which are indicated by color bars and are organized as follows: M (blue, male testes, male carcass), Fc (pink, NBF carcass, and multiple timepoints PBM: 12hr, 24hr, 36hr, 48hr, 60hr, and 72hr), Ov (orange, NBF ovaries, and multiple ovarian timepoints PBM: 12hr, 24hr, 36hr, 48hr, 60hr, and 72hr), D (tan, diapause at multiple timepoints: 0-1wk, 1-2wk, 2-3wk, and 3-4wk), Emb (embryo at multiple timepoints: 0-1hr, 0-2hr, 2-4hr, 4-8hr, 8-12hr, 12-16hr, 16-20hr, 20hr-24hr, 24-28hr, 28-32hr, 32-36hr, 36-40hr, 40-44hr, 44-48hr, 48-52hr, 52-56hr, 56-60hr, 60-64hr, 64-68hr, and 68-72hr embryos), L (gray, larvae 1st, 2nd, 3rd, and 4th instar larvae stages), and P (yellow, pupae, early male and female, and late male and female pupae stages). **B)** Read mapping analysis of *Ae. albopictus* developmental samples against the C6/36 cell line assembly (canu_80X_arrow2.2) and AalbF2 genome assemblies. The distribution reflects percentage of fragments mapped to too many loci (maroon), fragments mapped to multiple loci (blue), uniquely mapped fragments (dark blue). There is a significant reduction of duplication in AalbF2 genome assembly compared to the C6/36 cell line (Canu_80X_arrow2.2) genome. More transcripts fell under the uniquely mapped category in the AalbF2 genome.

The analyses of gene expression profiles across all developmental time points showed that the number of expressed genes (Transcripts Per Million ≥1) gradually increases through embryogenesis, reaching its highest peak at 68-72 hrs (Database S8). As previously observed, there is an increase in the number of expressed genes during the early pupal stages and the male germline expresses the highest number of genes among all samples (*56*). After a blood meal, female mosquitoes undergo a series of physiological changes to support oogenesis. In PBM ovaries, the number of genes expressed in the female germline changes dramatically from 12 hrs to 36 hrs.

Pairwise correlation analysis revealed that almost every developmental stage is most highly correlated with its adjacent stage and is very similar to what was previously found (Fig. S6) (*56*). To visualize the various patterns of gene expression and the relationships between the samples, hierarchical clustering and Principal Component Analyses was performed (Fig. S6).

Based on these analyses, embryos, PBM ovaries, pupae, larvae, and PBM female carcass samples tend to cluster closer together which is expected since their gene expression profiles are similar as these are developmentally related samples. Two notable exceptions include the male testes and early embryos (0-1 hrs, 0-4 hrs, and 4-8 hrs), likely due to transcripts related to the maternal-to-zygotic transition (Fig. S6). The male testes sample clusters away from all other samples, reflecting a distinguishing difference between this sample as compared to other samples sequenced (Fig. S6).

### Conclusions

AalbF2 and its associated gene set, databases of nrEVEs, miRNAs and piRNA clusters are collective resources that will enable great advances in *Ae. albopictus* biology. Additionally, we developed the first physical map of *Ae. albopictus*, which consists of fifty DNA markers that cover the largest genomic scaffolds, rDNA, PPO gene clusters, and the largest viral integration in the genome. Overall, FISH data were consistent with the assembled genome, confirming its large-scale structural accuracy. Combining *in situ* and bioinformatic approaches, we anchored to the *Ae. albopictus* chromosomes 58 scaffolds, whose length sum makes 75% of the genome. Analyses of mitotic chromosomes also showed that the *Ae. albopictus* chromosomes are slightly longer than *Ae. aegypti* ones, which is consistent with cytofluorimetry results.

Small RNA analyses identified 121 miRNAs including 26 novel miRNAs, some of which are strongly induced upon blood-feeding, suggesting important functions for these miRNAs in reproduction and development. piRNA cluster annotation has provided a high confidence set of piRNA clusters, setting the stage for their inactivation or modification to understand their functions and to explore avenues to exploit them to prevent arbovirus transmission. Moreover, the strong enrichment of newly annotated nrEVE sequences in piRNA clusters provides fuel for the hypothesis that they may provide a potential inherited antiviral defense system (*11*, *13*, *22*). Curation of immunity genes annotation, among the predicted 26,856 protein-coding sequences and the M locus will unable insights into the immunity pathways that contribute to *Ae. albopictus* vector competence and provide venues for novel genetic-based strategies of control, including those for population suppression based on gene-drive systems creating male-biased populations (*57*). The developmental transcriptome analysis described here demonstrates that the new genome assembly has produced a significantly more complete gene set with less gene duplications as compared to the previously available genome. The quantification data across developmental timepoints and multiple tissues will provide the community with an invaluable resource for further exploration of *Ae. albopictus* biology.

### Methods Summary

We first used a cytofluorimetric approach to estimate the size of the *Ae. albopictus* genome across geographic populations. Then, starting from 2 pools of 40 mosquitoes of an inbred strain derived from the reference Foshan strain, we produced 82 Gb of PacBio and 135 Gb of Hi-C sequence. Initial contigs were generated using Canu v1.7.1, followed by consensus polishing and haplotype purging that resulted in a primary (AalbF2) and an alternative assembly. The primary contigs were then scaffolded by SALSA v2.0 using the Hi-C data. Physical mapping by *in situ* hybridization of fifty DNA markers, followed by bioinformatic-based comparison to the *Ae. aegypti* genome assembly (AaegL5), were used to anchor the *Ae. albopictus* genome sequence to the chromosomes. The completeness and quality of AalbF2 were comparatively tested with respect to both AaegL5 and AloF1 using BUSCO (*6*), Barnap (https://github.com/tseemann/barrnap) and alignment parameters of published genomic and transcriptomic data. The AalbF2 assembly was deposited at NCBI in June 2019 and annotated using the NCBI RefSeq Eukaryotic gene annotation pipeline (https://www.ncbi.nlm.nih.gov/genome/annotation_euk/process/). Annotation Release 102, named Aalbo_primary.1 (ID: GCF_006496715.1), was made public in August 2019. The genome assembly was used for in-depth analyses of several features, including manual annotation of immunity genes, nrEVEs and estimates of linkage-disequilibrium and population differentiation using population genetic approaches on WGS data from two wild populations. Small RNAseq libraries were further generated from carcasses and ovaries of both sugar- and blood-fed mosquitoes and used to annotate miRNAs and piRNA clusters and study changes in their expression profile following a blood-meal. Finally, an atlas of the *Ae. albopictus* expressional profile was built from an extensive panel of 47 independent transcriptome datasets and was then used to study gene co-expression networks during mosquito development and following blood-meal. Details of methodological procedures and resources are described in the supplementary material.

## Acknowledgments

We would like to thank Francesca Scolari and Patrizia Chiari for mosquito maintenance.

## Author contributions

**conceptualization, resources and whole-genome sequencing**: M. Bonizzoni and J. Powell; **assembly and Hi-C-data-based scaffolding**: S. Koren, J. Ghurye, A. Rhie, A.M. Phillippy; **whole-genome assembly quality control and analyses**: U. Palatini, E. Pischedda, M. Bonizzoni, F. Krsticevic, P.A. Papathanos; **physical mapping**: R.A Masri, A. Sharma, D.A. Karagodin, J. Jenrette, M. Sharakova; **cytofluorimetric-based genome size analyses**: J.S. Johnston; **small-RNA seq libraries and sequencing**: M. Marconcini, A. Failloux; **nrEVEs annotation**: U. Palatini, E. Pischedda; **piRNA clusters and miRNAs annotation**: R. Halbach, P. Miesen, R.P. Van Rij; **genome annotation**: F. F. Thibaud-Nissen; **immunity genes**: F. Krsticevic, P.A. Papathanos; **M locus**: J. Biedler, Z. Tu; **population genomics**: L.V. Cosme, J.E. Crawford, A. Caccone; **developmental transcriptome**: I. Antoshechkin; S. Gamez; O.S. Akbari; **Figures**: U. Palatini, R.A. Masri, M.V. Sharakova, R. Halbach, P. Miesen, R.P. Van Rij, J. Biedler, Z. Tu, F. Krstivevic, P.A. Papathanos, L.V. Cosme, J.E. Crawford, A. Caccone, S. Gamez; O.S. Akbari; **manuscript writing team:** M. Bonizzoni, U. Palatini, J. Powell, P.A. Papathanos, M.V. Sharakova, R. Halbach, P. Miesen, R.P. Van Rij, J. Tu, A. Caccone, S. Gamez; O.S. Akbari; **Competing interests:** authors declare no competing interests. J. E. Crawford is employed by Verily Life Sciences LLC and has no competing interests; S. Koren has received travel compensation to present at Pacific Biosciences meetings. **Data and materials availability:** main text or the supplementary materials include all data and information on data accessibility.

